# Coupling renewable energy infrastructure planning and biodiversity conservation: a modeling-based framework

**DOI:** 10.1101/2024.08.30.610451

**Authors:** Jérémy S.P. Froidevaux, Isabelle Le Viol, Kévin Barré, Yves Bas, Christian Kerbiriou

## Abstract

Reconciling renewable energy planning and biodiversity conservation is urgently needed to address the interconnected crises of climate change and biodiversity loss. However, current strategies to avoid or limit the negative effects of renewable energy on biodiversity still hold major limitations during the planning process that could be overcome with modeling approaches. Here we propose a new applied modeling-based framework aiming to determine potential threats posed by proposed or built projects to biodiversity. By capitalizing on large-scale standardized citizen science biodiversity data to create reference scales of biodiversity levels, this approach aims to better inform the Ecological Impact Assessment (EIA) process at different stages pre- and post-project construction. We demonstrate the practical application of the framework using bat and onshore wind energy development in France as a case study. We reveal that current approaches in renewable energy planning failed to identify sites of biodiversity significance with >90% of wind turbines approved for construction to be placed in sites of high significance for bats. The risks posed by future wind turbines to bats concern all taxa (that are all protected in the European Union), including species with higher collision risks. We highlight how the proposed modeling-based framework could contribute to a more objective evaluation of pre- and post-construction impacts on biodiversity and become a prevalent component of the EIA decision-making. Its implementation could lead to a more biodiversity-friendly renewable energy planning in accordance with the world-leading target to halt biodiversity decline by 2030.

## 1. Introduction

It is now well recognized that climate change and biodiversity loss are fundamentally intertwined and that both crises should be addressed collectively [1-3]. While in many economic sectors actions to combat climate change may also contribute to biodiversity conservation and restoration and vice-versa (e.g. agri-environment-climate measures in European farmland) [4], climate change mitigation measures in the energy sector such as the development of renewable energy sources are not free from negative impacts on biodiversity [5-7]. The urgent need of moving away from fossil fuel to carbon-free energy production for reducing global CO2 emission therefore conflicts with the overreaching global goal of halting and reversing biodiversity loss [8], thus leading to the so called “green-green dilemma” [7, 9] between two key Sustainable Development Goals adopted by the United Nations in 2015. Nevertheless, strategic renewable energy planning, siting and mitigation measures could avoid or limit the negative impacts renewable energy have on biodiversity during their construction, operation, and decommissioning, and represent a compelling opportunity in the energy sector for reconciling climate action and biodiversity conservation [10]. Thus, in many countries worldwide renewable energy projects – like other development projects (e.g. road and railway construction) – fall within the mitigation hierarchy, a decision-making framework for mitigating biodiversity loss from development by sequentially avoiding, reducing and a last resort offsetting any potential impacts with the ambition towards no-net-loss of biodiversity [11, 12]. By identifying and evaluating the potential impacts of a proposed project on biodiversity and proposing ways to avoid/mitigate/offset these impacts, the ecological impact assessment (EIA) represents a cornerstone of the mitigation hierarchy [13].

In the European Union, the Directive 2011/92/EU outlines the process of EIA which ensures that projects that are likely to have a significant impact on biodiversity are assessed prior to their authorization. In brief, following the screening and scoping stages this typically includes (i) a pre-selection of potential sites based on spatialization and mapping of the impacts, (ii) field-based ecological surveys of protected species and habitats (Habitats Directive 92/43/EEC and Birds Directive 79/409/EEC) on the pre-selected sites to document the EIA, and (iii) an impact analysis in which potential impacts and their strength are identified and quantified and mitigation measures are proposed. In some cases (for instance in wind turbine project development), EIA process continues after the construction of the proposed projects as some of them can be subject to post-construction impact assessment to evaluate whether the predicted impacts have occurred and if the mitigation measures were effective. However, while the EIA process is regulatory, it holds major limitations that can lead to serious errors during decision-making processes [14]. For instance, the identification of potential sites using spatialization and mapping tools most often rely on incomplete (e.g. data collected with inadequate sampling effort) and biased biodiversity data (including taxonomic, temporal and spatial biases). While regulatory field-based ecological surveys allow to ground-check this first assessment, their implementations are far from being optimal [14]. First, these surveys are constrained by financial resources which limit the sampling effort allocated [15, 16]. This is especially problematic since the impact analysis mainly arises from biodiversity data collected during the surveys and very rarely considers other forms of biodiversity such as (i) the undetected biodiversity (i.e. species present in the site but not detected during the survey) [17], (ii) the expected biodiversity (i.e. species capable of coexisting in the study site at the present time but due to stochastic events are absent from the site during the surveys; for instance the dark biodiversity [18]), (iii) the biodiversity potential (through predictions) and (iv) the possible changes in biodiversity (through scenario analyses). Second, the evaluation of a site’s significance in terms of biodiversity can be restricted by the limitations of available external references, which frequently depend on heterogeneous and non-standardized data. Overall, the whole EIA process crucially suffers through its different stages from the lack of standardization for biodiversity data interpretation and contextualization, resulting in subjective evaluation of pre- and post-construction impacts [19, 20].

One way to overcome these last issues that are currently inherent to EIA is to create reference scales of biodiversity levels [20] that can be defined at different spatial (from local to national and international levels) and temporal (dynamic over time) scales. To that end, large-scale ecological models of how species distributions and abundances vary over space and time (i.e. abundance-based species distribution models) could represent an effective tool. Species distribution models (SDM) are well-established in the scientific literature [21, 22] and have already proved to be of great asset for assessing the effectiveness of the EIA procedure [23, 24] and guiding conservation planning [25-28]. Nevertheless, building reliable reference scales of biodiversity levels using SDM requires a massive amount of biodiversity data collected in a standardized way through space and time. Although there are extensive databases available containing big biodiversity data (e.g. Global Biodiversity Information Facility database), a considerable amount of the data in these databases is obtained opportunistically with unknown collection processes, and is subject to observer’s bias, producing biases such as over-sampling of flagship species, favorable areas or habitats [29, 30]. These biases are difficult to correct or involve complex analysis [31, 32], and together with the lack of sampling effort quantification make the data unsuitable for the intended purpose. In contrast, standardized biodiversity monitoring schemes – especially those that are intended to detect large-scale spatiotemporal trends of abundance and distribution – could be an important source of high-quality data on species abundance. These schemes have been developed in many countries worldwide and for many taxa, including insects (e.g. [33]), birds (e.g.[34]) and mammals (e.g. [35]). They have the advantage of being based on standardized protocols with a fixed sampling effort and allows for comparison among sites without relying on assumptions about observer site or species preferences or adjusting for varying sampling efforts [36]. Depending on their goals, these monitoring schemes aim to represent the current national distribution of habitats and biogeographic context. While financial and logistic constraints may sometimes limit their coverage, biodiversity monitoring schemes that are based on citizen science and adhering to standardized protocols have the potential to provide extensive, standardized biodiversity data [37, 38].

Here, the aim of the study was to demonstrate how reference scales of biodiversity levels derived from large-scale standardized citizen science biodiversity monitoring programs could be used to inform decision-making processes at various stages of the mitigation hierarchy process. We developed an applied modeling-based framework to determine (i) prior to field-based ecological surveys whether projects proposed for development are in areas of biodiversity conservation significance and (ii) post construction whether the project complies with the mitigation hierarchy framework. The modeling framework includes five main steps (Figure 1): (i) identifying large-scale citizen-science programs and determining relevant predictors (e.g. environmental and bioclimatic variables) known to shape the spatial distribution of the target species potentially occurring in the study area; (ii) modeling species abundance using biodiversity data from large-scale citizen science programs in relation to previously identified relevant predictors; (iii) predicting species abundance at random points to build a standardized reference scale of species abundance; (iv) predicting species abundance at the proposed/built sites and comparing the prediction to the standardized reference scale of species abundance to evaluate potential risks posed by the proposed/built projects; (v) identifying sites of potential biodiversity significance threatened by the proposed/built projects. We developed and tested the modeling framework using bats and onshore wind energy development in France as a case study. In fact, the negative impacts of onshore wind turbines on bats are well documented and include both fatalities [39] and losses of habitat use due to avoidance [40, 41]. France represents an ideal case study as (i) it is one of the largest wind energy contributors in the EU with a pace of wind turbine installation currently undergoing rapid acceleration, and (ii) it holds the largest national-scale standardized citizen-science bat monitoring program in Europe. According to the mitigation hierarchy established in the Article 6 of the EU Habitats Directive 92/43/EEC, we expected that wind turbines approved by local environmental planning authorities for construction (i.e. projects that have undergone an EIA) would be localized in areas of low bat activity levels for all taxa (as bat species are listed in Annex IV of the EU Habitats Directive) and especially for taxa with higher collision risks. We provide through this case study a concrete illustration on how the modeling framework can be applied in EIA to inform the decision-making process.

**Figure 1.**
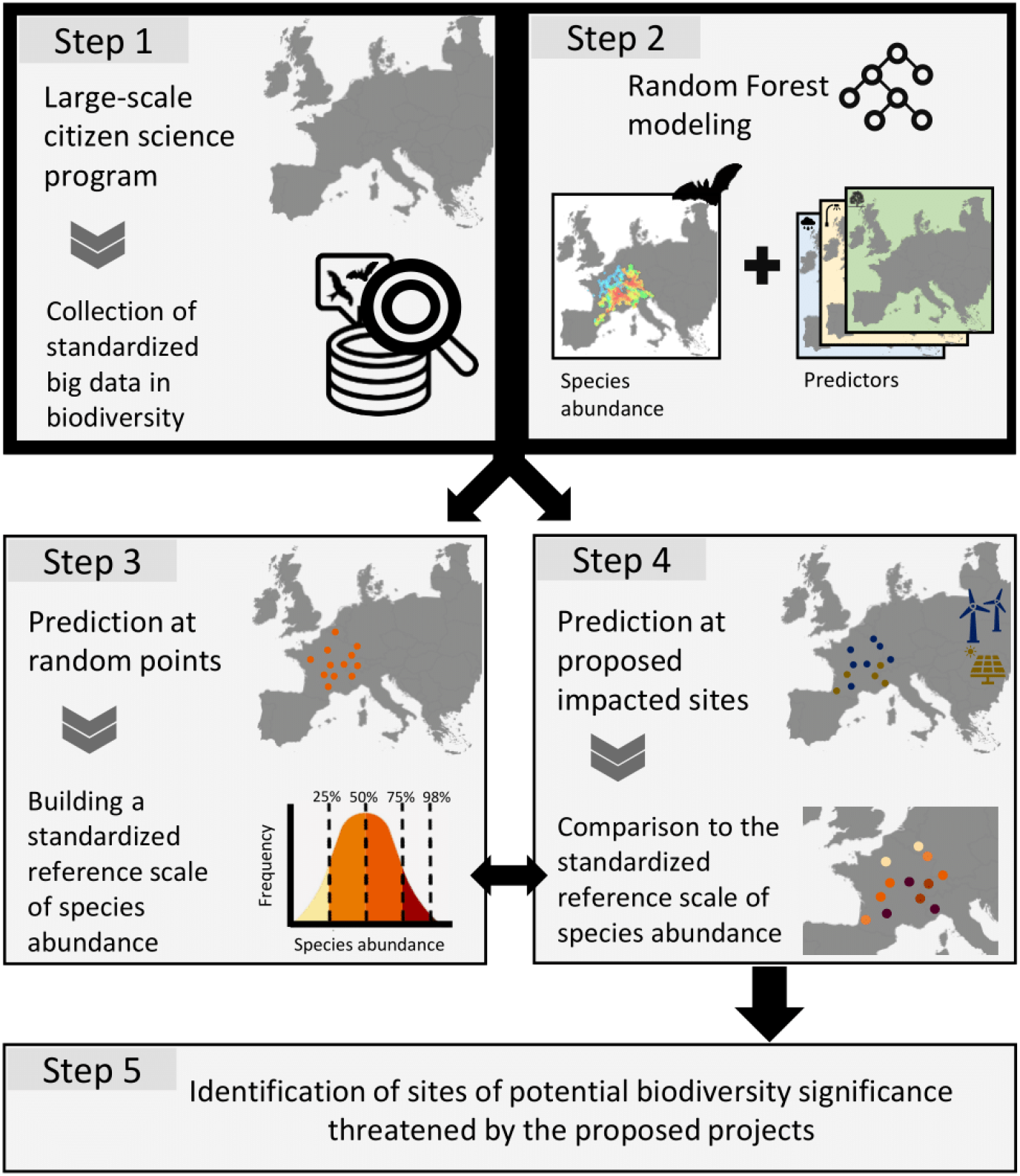
Conceptual modeling-based framework proposed to be implemented within the ecological impact assessment process.

## 2. Material and methods

### 2.1. Study areas

The case study focused on two areas in France with contrasting past and current wind farms development, namely the region Bourgogne-Franche-Comté (BFC, eastern France, 47°12’ N, 4°57’ E, 47,783 km^2^, altitude: 52 to 1,495 m a.s.l.) and the regions Bretagne and Pays de la Loire (BPL, western France, 48°12’ N, 2°55 W, 59,290 km^2^, altitude: 0 to 416 m a.s.l.). The BFC region is mainly covered by forests (36%) and grassland (32%) while two-thirds of the BPL area consist of agricultural lands – with 36% of arable lands and 31% of grassland. By mid-2020, a total of 388 and 1,105 wind turbines were operational in BFC and BPL, respectively, and 233 and 766 were approved by local environmental planning authorities for construction in BFC and BPL, respectively.

### 2.2. Modeling framework for assessing potential ecological impact: case study with bats and wind turbines

#### 2.2.1. Step 1a: Using large-scale database and evaluation of their representativeness

Here we used data from the French national-scale citizen-science bat monitoring program “Vigie-Chiro” [42]. We retrieved bat activity data from surveys conducted between 2015 and 2020 following the stationary points protocol in which trained volunteers acoustically sampled bats during at least one full night when weather conditions are optimal for bats to forage (see [43-45] for more details). Trigger settings were standardized among detectors to minimize heterogeneity in detectability (https://www.vigienature.fr/fr/page/participer-vigie-chiro). Species identification was conducted using Tadarida software [46] which automatically detects and extracts sound parameters of recorded echolocation calls within a 5-sec file (i.e. a bat pass) and classifies them into bat taxa with an associated confidence index. We followed the method proposed by Barré, Le Viol [47] to account for potential automated identification errors, i.e. we used the confidence index to retain two separate datasets: (i) one dataset of bat passes with maximum error risk tolerance of 10%; and (ii) another dataset of bat passes with maximum error risk tolerance of 50%. The first threshold is cautious, aiming to minimize false positives, while the second one is less conservative, allowing for a larger amount of data to be retained. Bat activity per night corresponded to the sum of bat passes recorded.

We restricted our selection to sites (i) sampled between May and October (i.e. period of highest bat activity), (ii) with microphones placed at <5 m height, (iii) located away (>200 m) from known roosts, and (iv) at lower altitude (<800 m a.s.l.) to avoid excessive heterogeneity due to mountainous environmental characteristics. In total, 711 sites corresponding to 1,186 detector-nights and 643 sites corresponding to 880 detector-nights were retained in BPL and BFC, respectively (Supplementary Material S1). Although the sites sampled are nationally representative [44], they were chosen through a participatory process, which may result in spatial distribution heterogeneities at the local or regional level. Therefore, it is crucial to evaluate the potential impact of this heterogeneity on representativeness and to develop appropriate strategies to address any spatial structure issues. Regarding the representativeness, the amount of major land-cover classes around the sampling sites were representative of the two study areas, except for BFC where deciduous forests were slightly over-sampled (Supplementary Material S2). Similarly, gradients of environmental variables and anthropogenic pressures around the sampling sites matched the gradients observed within the two study areas (Supplementary Material S2). Furthermore, there was no confounding effect between habitat type surveyed and detector type used (Supplementary Material S2). Because sampling sites (especially in BPL) were clustered, we developed in step 2 (see section 2.2.3) models using the full dataset and models using a subset of the dataset containing only sites that were located >500 m away from each other to account for potential spatial structure issues. We conducted the analysis on 12 bat species or species groups (due to current challenges in species identification using acoustic monitoring) recorded in the two study areas (Table 1).

**Table 1.** Summary of percentage of occurrence and mean bat activity per night across sites of the 12 bat taxa studied in the two study areas. Values are given considering an identification of bat passes with a maximum error risk tolerance of 50%.

| Taxa | Abbreviation | Bourgogne-Franche-Comté (N = 643) |  | Bretagne - Pays de la Loire (N = 711) |  |
| --- | --- | --- | --- | --- | --- |
|  |  | % of occurrence | Mean bat activity | % of occurrence | Mean bat activity |
| *Barbastella barbastellus* | B.bar | 68 | 12 | 74 | 17 |
| *Eptesicus serotinus* | E.ser | 78 | 45 | 65 | 18 |
| *Myotis nattereri* | M.nat | 65 | 7 | 68 | 14 |
| *Myotis* spp. (excluding *M. nattereri*) | Myo.spp | 84 | 73 | 78 | 50 |
| *Nyctalus leisleri* | N.lei | 72 | 21 | 52 | 7 |
| *N. noctula* | N.noc | 40 | 6 | 21 | 10 |
| *Pipistrellus pipistrellus* | P.pip | 99 | 709 | 100 | 786 |
| *P. nathusii/kuhlii* | P.natkuh | 79 | 82 | 95 | 176 |
| *P. pygmaeus/Miniopterus schreibersii* | P.pygM.sch | 33 | 3 | 23 | 1 |
| *Plecotus* spp. | Plec.spp | 39 | 2 | 63 | 5 |
| *Rhinolophus ferrumequinum* | R.fer | 23 | 2 | 28 | 6 |
| *R. hipposideros* | R.hip | 47 | 5 | 29 | 8 |

#### 2.2.2. Step 1b: determining relevant predictors

We collected 40 predictor variables that are relevant for predicting bat activity in human-modified landscapes [48-51]. These variables were contiguously available at high resolution for the two study areas (see Supplementary Material S3 for more details on data sources, resolution and temporal coverage) and include (i) two topographic predictors (altitude and slope), (ii) eight environmental and land-uses variables (the proportion and Euclidean distance to deciduous forests and linear small woody features, the density of rivers and distance to the nearest freshwater body, the edge density and Shannon’s diversity index of major land-cover classes), (iii) 11 predictors related to anthropogenic stressors (artificial night-time light brightness, amount and distance to urban areas and croplands, the density and distance to major roads and operational wind turbines, human population density, quietness suitability index), and (iv) 19 bioclimatic predictors. For BPL which is bordered by the Atlantic Ocean, we also derived the distance to the coastline.

Bats are mobile taxa that respond to environmental variables and anthropogenic stressors at different spatial scales, from local to landscape scales [48]. We therefore implemented a multi-scale approach and derived area-based variables across ten spatial scales. We used ArcGIS Desktop v10 (ESRI, Redlands, CA, USA) to create ten buffers of 0.05, 0.10, 0.25, 0.50, 1.00, 2.00, 3.00, 4.00, 5.00, and 10.00 km radius around each sampling site. The large scales were selected considering the mean and maximum daily foraging movement of European bat species [52] whereas the small ones allow us to have a fine-scale description of the near environment of the sampling sites. The density of operational wind turbines was calculated for the six largest spatial scales only as most sites (>75%) were located >1 km away from wind farms. Given the recent increases in wind turbines installation between 2015 and 2020 in France, we considered the two predictors related to wind turbines as dynamic [53], i.e. we calculated the density and distance to wind turbines of a given site using only wind turbines that were operational at the time of the bat survey.

#### 2.2.3. Step 2: modeling species abundance

We used Random Forest (RF; [54]) implemented in the R package randomForest [55] to model bat activity in relation to topographic, environmental, anthropogenic and bioclimatic predictors variables in each study area (BFC and BPL). RF models were parameterized with the recommended default values (ntrees = 500, cutoff = 1/k = 1/3). We added temperature at night, Julian day and the site coordinates in the final list of predictors to consider spatiotemporal variation in bat activity. We used participant ID and site ID as strata to account for the stratified structure of the citizen-science data. While RF can operate with large numbers of variables and is largely insensitive to multicollinearity, it is recommended to proceed to variable selection to improve overall model performance [56]. We therefore conducted a variable selection procedure using the package vsurf [57] and retained the smaller set of variables sufficient for prediction purposes. In this process, predictor variables were assessed for their individual impact on model performance and the resulting list of selected predictors was refined by eliminating redundancy. Final models that included the selected variables explained higher variance (here referred to as a measure of how well the out-of-bag (OOB) predictions capture the variability of the target variables in the training set) than full models (Supplementary Material S4) and were therefore retained to proceed to model performance evaluation and prediction. We assessed model performance using the Normalized Root Mean Square Error statistic (NRMSE) derived from a fivefold cross□validation procedure. NRMSE measures the divergence of the predictions generated by the models (using a training set) from observations of a test set. We iterated 500 times the fivefold cross□validations to calculate NRMSE, with a training set consisting of 80% of the observations randomly selected at each iteration and a testing set corresponding to the remaining 20% observations. We provide an Overview, Data, Model, Assessment and Prediction (ODMAP) [58] as part of the Supplementary Material S5.

Using citizen-generated acoustic data to model bat activity comes with some challenges that need to be addressed during the modeling process. First, we accounted for uncertainties in bat identification by modeling bat activity using two separate datasets (hereafter referred to as “acoustic datasets”) having different error risk tolerance thresholds in acoustic identification (10 vs 50%) [47]. Second, because some Vigie-Chiro sites were spatially clustered (especially in BPL) - thus potentially leading to an overestimation of model performance - we also ran the final models using two datasets (hereafter referred to as “spatial datasets”) that included either all the sites or only spatially independent ones, i.e. a subset of the full dataset containing only sites that were located >500 m away from each other (N_BPL_ = 473, N_BFC_ = 546).

#### 2.2.4. Step 3: building a standardized reference scale of species abundance

We built a reference scale of bat activity for each study area in three steps. First, within each area, we randomly selected the same number of random points as there were wind turbines approved for construction (i.e. 233 in BFC and 766 in BPL). We employed a random stratified approach: random points were located (i) within a 50 km radius from wind turbines approved for construction, for encompassing areas with similar environmental and bioclimatic conditions, (ii) >150 m from each other (with >98% of points located > 500 m); and (iii) outside urban and protected areas because wind turbines sitting in such area is not permitted. We were not able to exclude other restricted areas (e.g. zones of aeronautical easements) as GIS layers were not publicly available. Second, we used the final RF models to predict species-specific bat activity at these random points. Third, we built the reference scale of bat activity using the ordered value of bat activity predicted at random points. We used percentile threshold [20, 59] with the following five categories: (i) low activity: 0-25th percentiles; (ii) medium-low: 25-50th percentiles, (iii) medium-high: 50-75th percentiles, (iv) high: 75-98th percentiles, and (v) extremely high: 98-100th percentiles.

#### 2.2.5. Steps 4 and 5: predicting species abundance at proposed sites and assessing potential risks posed by wind turbines to bats

To provide a concrete illustration through this case study on how the final steps of the modeling-based framework can be applied in EIA to inform the decision-making process, we assessed whether wind turbines approved by local environmental planning authorities for construction (i.e. projects that undergone an EIA) were effectively in areas of low bat activity levels. Indeed, we expected that the approved wind turbines will be sited away from important foraging and commuting habitats for bats since bats are strictly protected in the European Union (Habitats Directive of the European Union 92/43/EEC) and are considered in the mitigation hierarchy process during wind turbines planning since several years in France. To do so, we used the final RF models developed in step 2 to predict species-specific bat activity at the wind turbines approved by local environmental planning authorities for construction and then compared the predictions obtained to the standardized reference scale of bat activity previously built in step 3 for each taxon at the regional level. This comparison allowed us to quantify potential risks posed to bats by the approved wind turbines.

We finally evaluated species-specific patterns in risks posed by wind turbines to bats as the type of impacts of wind turbines on bats are largely species-specific, with mortality events by collision mainly affecting high-flying species [60]. We tested whether future wind turbines that will be placed in areas with high bat activity would regard only bats that are not subject to high collision risks, as one would expect if the mitigation hierarchy process was successful in avoiding direct impacts. To do so, we first retrieved for each species independently the proportion of wind turbines that will be built in areas of high and extremely high activity levels. Then, we tested the relationship between the proportion obtained for each species in relation to their collision susceptibility index (in a logarithmic scale due to the large spread of values) using beta regression models [61]. Collision susceptibility index was obtained from Roemer, Disca [60]. Models were checked using the performance package [62].

## 3. Results and discussion

### 3.1. Overview of random forest models

The modeling-based framework was developed and tested using 12 bat taxa which were monitored through the French national-scale citizen-science bat monitoring program “Vigie-Chiro” in two distinct areas in France – Bourgogne-Franche-Comté (BFC) and Bretagne-Pays de la Loire (BPL). We built 96 RF models, i.e. one per taxa and per area and considering (i) two separate acoustic datasets having different error risk tolerance thresholds in acoustic identification (10 vs 50%), and (ii) two spatial datasets that included either all the sites or only spatially independent ones (full vs subset dataset). The predictive performances of random forest models were overall satisfying with most NRMSE values below 20% [63], even though predictive performance varied to some extent with respect to species, area and spatial dataset considered (Supplementary Material S6). Given the overall predictive performance of the RF models developed and for sake of clarity and concision, results of subsequent analyses are only reported for the 50% full dataset.

The variable selection procedure (step 2) led to different combinations of predictor variables for the 12 taxa in the two study areas (Figure 2). The top ranked predictors retained in >25% of models included climate variables, quietness suitability index and amount of cropland at different spatial scales, distance to water, and Julian day. Other key predictors shared at least between four taxa included the amount of deciduous forest, small woody features, urban areas as well as river density at several spatial scales, and distance to cropland and wind turbine (Figure 2). Overall, environmental variables retained in the final models fit with our expectations regarding bat ecology and their responses to anthropogenic stressors [49]. For instance, our results are in line with Azam, Le Viol [64] who demonstrated that the amount of cropland in the landscape was the main factor negatively affecting four common bat species in France. Similarly, several studies demonstrated that proximity to resources such as freshwater sites is a key driver of bat activity [65-67]. The variable selection process also emphasized the importance of considering the quietness suitability index when modeling bat activity, which is consistent with increasing field-based evidence regarding the effects of anthropogenic noise on bats [68, 69]. In addition to noise, quietness suitability index may encompass other urban-related stressors (e.g. major road density, artificial night-time light brightness, impervious surface) making it a crucial predictor to consider when modeling bat activity.

**Figure 2.**
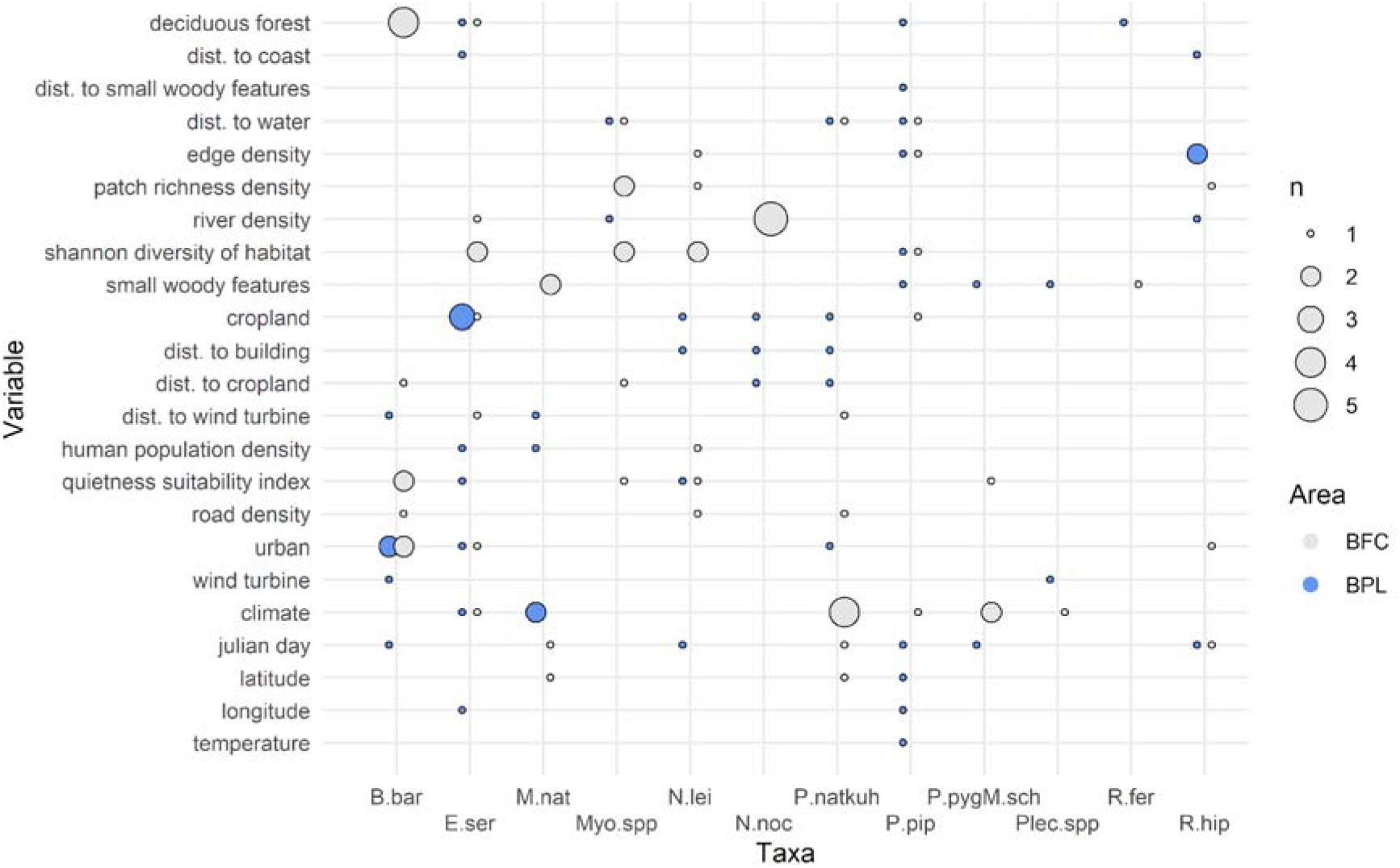
Summary of the variables retained in the final random forests built for 12 bat taxa in two study areas (BFC: Bourgogne-Franche-Comté, BPL: Bretagne-Pays de la Loire) using the 50% full dataset (see the Results section for more details). The presence of a dot at the intersection between a given variable and a given species means that the variable was retained in the final random forest model built for that species. The color of the dots indicates the two study areas (gray: BFC, blue: BPL). The dot size corresponds to the number of retained spatial scales for environmental variables and the number of selected climate variables for the climate variable.

The reliable predictive performance of models, along with consistent species responses corroborating findings in the literature, reinforce the notion that citizen science-based biodiversity monitoring schemes adhering to standardized protocols could provide high-quality big data [37, 38]. The use of standardized acoustic data collected through the national-scale citizen-science bat monitoring program Vigie-Chiro in combination with high-resolution climatic and environmental variables has proved to be valuable for modeling bat activity. The development of such models would not have been possible with other sources of data since standardized abundance/activity data on bats are scarce or not centralized and often limited in spatiotemporal extent, thus hampering large-scale modeling studies. Nevertheless, combining data from acoustic and roost monitoring programs would be required to adequately model the abundance of species that are difficult to detect acoustically. Overall, citizen-science bat monitoring program as implemented in many countries worldwide [42, 70-73] represents a rich source of big data that is expanding over time and that have the potential to better inform evidence-based management and conservation strategies.

### 3.2. Failure in current mitigation hierarchy process highlights the crucial need of adopting a modeling-based approach

The comparison of predicted species-specific bat activity at the wind turbines approved by local environmental planning authorities for construction (step 4) to the standardized reference scale of bat activity (step 3) highlighted that a considerable proportion of wind turbines will be placed in areas with high or extremely high bat activity levels (Figure 3a). While this proportion varies among species, we found that only less than 10% of wind turbines will be placed in areas where no high or extremely high bat activity levels of any taxon is expected. At the other side of the spectrum, more than 25% of wind turbines will be in areas of high and extremely high activity for one third of the bat assemblage (Figure 3b). When investigating in more details species-specific pattern, we found no significant relationship (BFC: *P* = 0.19, estimate ± SE = -0.06 ± 0.05; BPL: *P* = 0.34, estimate ± SE = -0.04 ± 0.05) between the percentage of wind turbines in areas of high and extremely high activity and the collision susceptibility index of the taxa that will be affected by the wind turbines (Figure 3c). In other words, wind turbines will be built in areas of high bat activity where even species with higher collision risks are expected to be very active. If the current mitigation hierarchy process was truly effective in avoiding the direct impacts of wind turbines on bats, we would have expected to observe a much lower proportion of wind turbines located in areas of high or extremely high bat activity for species with higher collision risks. Thus, our results corroborate those of Lintott, Richardson [14] who provided empirical evidence that ecological impact assessments fail to reduce risk of bat casualties at wind farms in the United Kingdom. Nevertheless, we acknowledge that information regarding the presence of curtailment – i.e. a reduction measure consisting of operational restriction of the turbines during high bat activity – were not provided. In addition, the more subtle effect of habitat loss due to wind turbine avoidance by some bat species [40, 41] were not considered in the analysis due to the lack of consideration of this type of impact in past and current EIA process [74].

**Figure 3.**
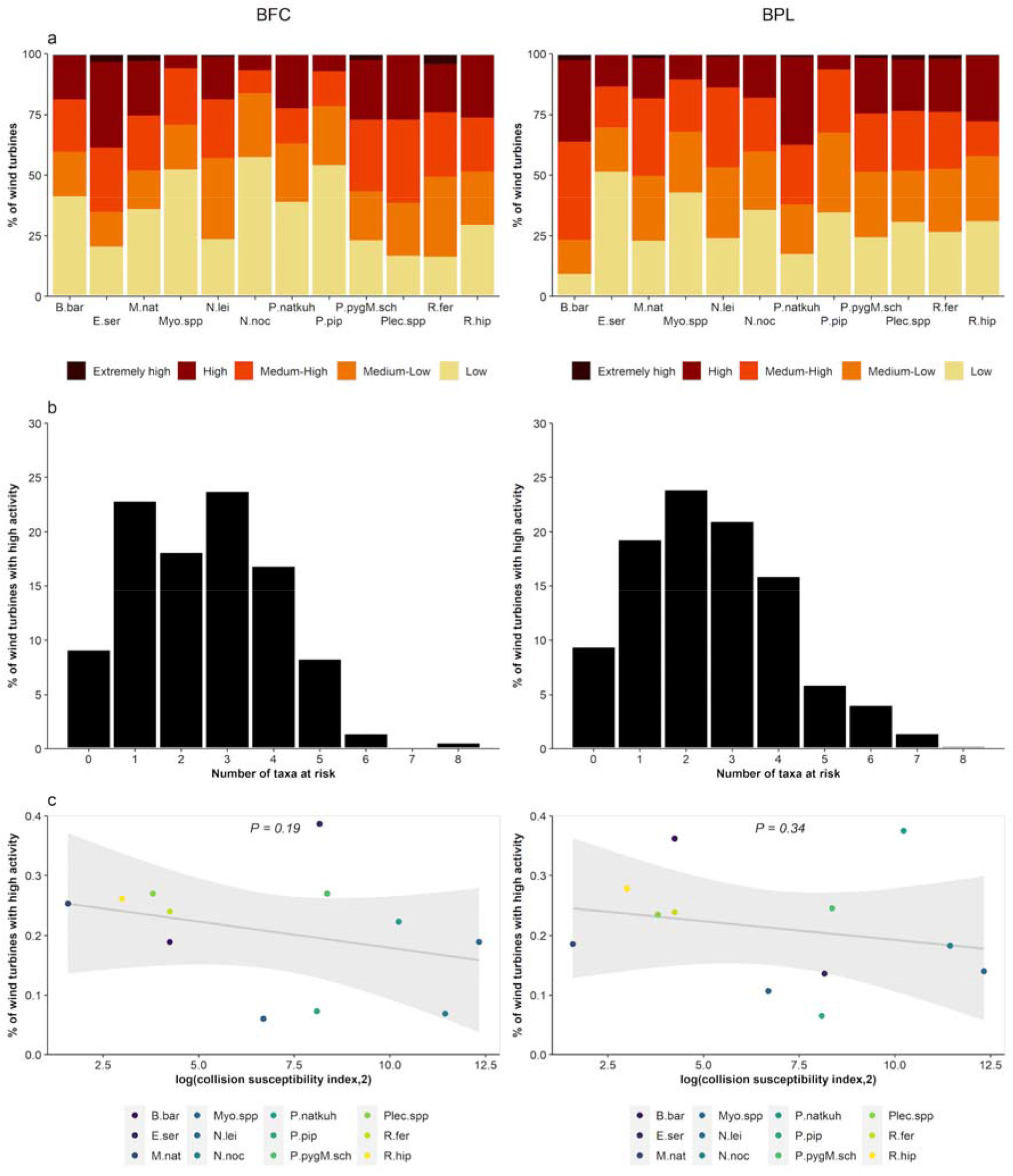
Potential risks posed by wind turbines approved by local environmental planning authorities for construction to bats in the two study areas (BFC: Bourgogne-Franche-Comté, BPL: Bretagne-Pays de la Loire). **(a)** Stacked bar plots depicting the percentage wind turbines that will be placed in areas of low, medium-low, medium-high, high, and extremely high levels of activity for each bat species. **(b)** Bar plots showing the percentage of wind turbines that will be sited in areas of high and extremely high bat activity in function to the number of taxa they will put at risk. **(c)** Relationship between the percentage of wind turbines with high and extremely high activity and the collision susceptibility index of the taxa that will be affected by the wind turbines.

This study case highlights the crucial need of adopting a modeling-based approach to derive robust reference scales of biodiversity levels that will allow to (i) assist during spatialization and mapping of the impacts prior to regulatory ecological ground surveys and determine whether the projects proposed for development are in area of biodiversity conservation significance, and (ii) assess post construction whether the projects comply with the mitigation hierarchy framework (Figure 4). For instance, the whole modeling-based framework could be implemented as a toolbox for stakeholders (e.g. ecological consultants, environmental planning authorities) to aid in predicting potential impacts of future projects on biodiversity. This approach could also help identifying infrastructures already sited in areas of predicted high species abundance and advice for targeted post-construction surveys and the implementation of mitigation measures for poor-sited ones (e.g. curtailment for wind turbines [75]). This approach is intended to be complementary, not substitutional, of the current EIA stages during renewable energy infrastructure planning. Indeed, while the modeling procedure allows to predict mean species abundance with regards to climate and environmental factors, regulatory field-based ecological surveys will always be required to capture site-specific variations in species abundance, which are influenced by the unique characteristics of the area.

**Figure 4.**
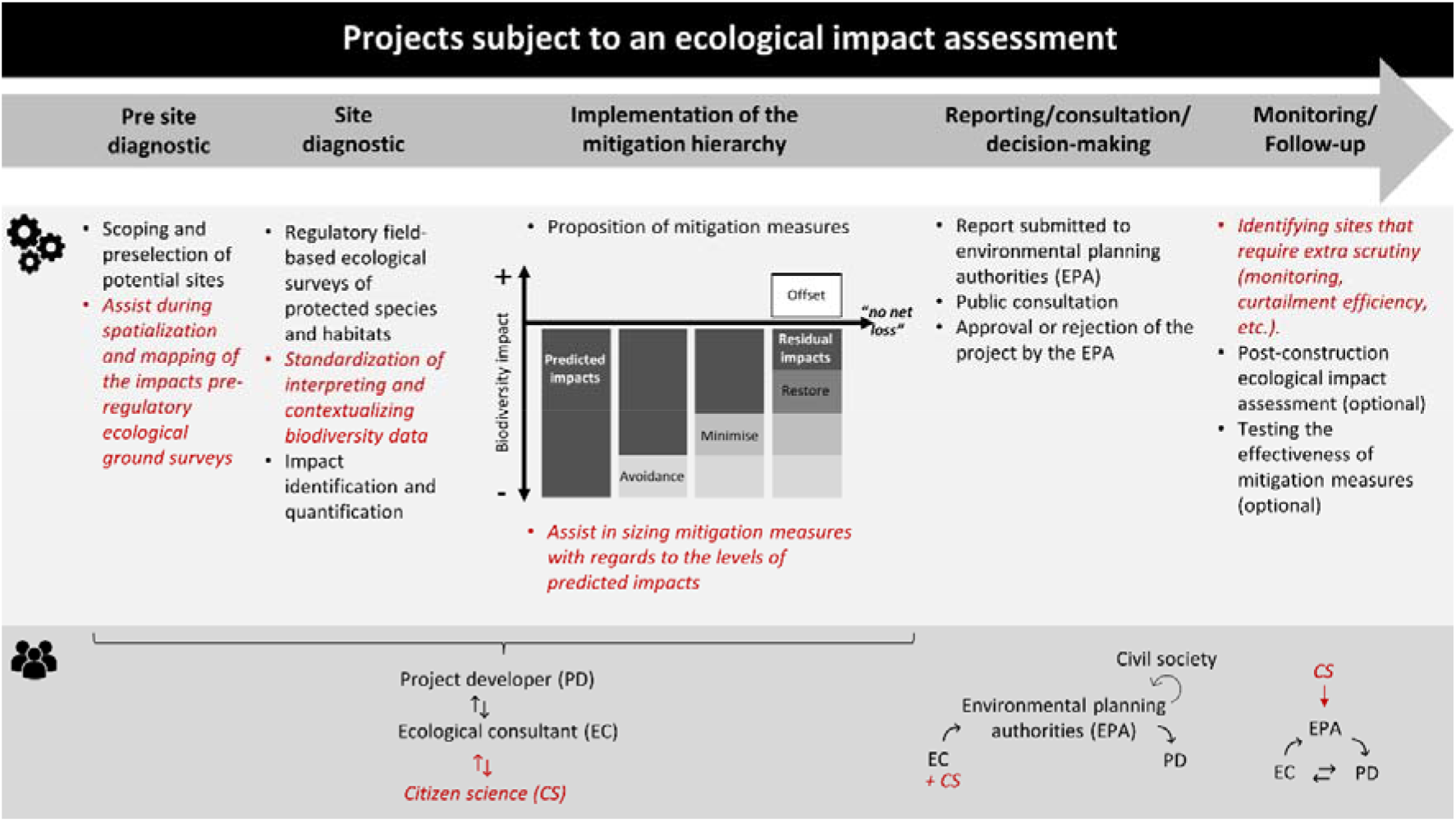
Potential contributions and adding values (in red) of integrating the proposed modeling-based framework in the ecological impact assessment process.

Finally, in light of the ongoing debate on whether important conservation areas are effective proxies for predicting the impact of renewable energy expansion on biodiversity [76, 77], the proposed modeling-based framework is well-suited to identify areas of biodiversity significance both inside and outside these priority conservation areas.

### 3.3. Towards contemporary, dynamic, and multi-season reference scales of biodiversity levels

Our method for determining the reference scales of biodiversity levels was similar to the contemporary reference state approach outlined in McNellie, Oliver [78]. This approach offers the advantages to (i) reevaluate and adjust the reference scales of biodiversity levels when additional data become available, and (ii) adapt the references scales in a context of rapid biodiversity change [79] principally caused by climate and land-use changes [80-82]. For instance, the reference scales of bat activity levels in France are intended to be dynamic to better consider spatial and demographic responses of bats to global changes [83, 84]. Nevertheless, we acknowledge that this approach solely facilitates the spatial identification of sites of potential biodiversity significance based on current references, without considering any potential past impacts. There is a trade-off between adapting the reference scales due to rapid biodiversity change in the Anthropocene and accounting for issues related to shifting baseline syndrome [85]. This trade-off should be considered depending on the targeted species and areas, as well as data availability.

Reference scales of biodiversity levels should also account for seasonal changes in species distribution and abundance which reflect key life-history events and seasonal species responses to climatic and environmental factors in both migratory and non-migratory species [86-88]. Given the increasing amount of within-year temporal biodiversity data in biodiversity monitoring programs and the increasing availability of climatic and environmental layers at high spatiotemporal scales [89], building robust multi-season reference scales of biodiversity levels will soon become an achievable target.

## 4. Conclusions

Using bats and onshore wind energy development in France as a case study, we demonstrated how reference scales of biodiversity levels modeled from large-scale standardized citizen science biodiversity monitoring programs could be effectively used to identify sites of potential biodiversity significance threatened by proposed or built projects. The implementation of the proposed modeling-based framework through this case study has allowed us to unveil that only less than 10% of wind turbines approved by local environmental planning authorities for construction will be placed in sites of little significance for bats. Furthermore, we found that the risks posed by future wind turbines to bats concern all taxa, including species with higher collision risks. Altogether, results of the case study suggest that the mitigation hierarchy process as currently implemented in France has failed to identify renewable energy projects with significant impacts on biodiversity, at least on bats. Adopting a modeling-based approach within the ecological impact assessment process to derive robust reference scales of biodiversity levels seems crucial to better assessing the impacts of a development project pre and post construction and comply with the mitigation hierarchy framework with the ambition of no-net-loss to biodiversity.

We illustrated in this study how reference scales of biodiversity levels allow to inform the mitigation hierarchy process through its different stages. In addition to assisting in spatialization and mapping of the impacts of pre-regulatory ecological ground surveys, the proposed modeling-based approach enables the standardization of interpreting and contextualizing biodiversity data. Thus, it contributes to a more objective evaluation of pre- and post-construction impacts on biodiversity, which ultimately could lead to a more biodiversity-friendly renewable energy planning in accordance with the world-leading target to halt biodiversity decline by 2030 [90, 91]. The modeling of reference scales of biodiversity levels requires a massive amount of biodiversity data collected in a standardized way and the lack of such data could therefore represent the main limitation of its relevance globally. Yet, large-scale standardized biodiversity monitoring schemes to track ecological changes over space and time have been – and still are – increasingly developed for a wide range of taxa in many countries worldwide. Thus, the implementation of this modeling-based framework in the mitigation hierarchy process has the potential to become a prevalent component of the EIA to better inform decision-making in renewable energy project development and beyond.

## Acknowledgments

We are grateful to all the volunteers of the Vigie-Chiro program. The success of such long-term, large-scale surveys relies entirely on their continuous involvement. We thank CC-IN2P3 and PCIA-MNHN for providing computing and storage facilities of acoustic data collected through Vigie-Chiro, and Didier Bas for his help in this process.

## Funding

JSP Froidevaux was funded by the SAD–Région Bretagne (SAD grant number: 19041) and the Leverhulme Trust through an early career fellowship (Award Reference: ECF-2020-571).

## CRediT authorship contribution statement

CK and ILV secured the funding from the SAD–Région Bretagne; JSPF, KB and CK conceived the modeling-based framework; JSPF analyzed the data and performed the case study; JSPF led the writing of the manuscript; All authors critically contributed to the drafts and gave their final approval for publication.

## Declaration of Generative AI and AI-assisted technologies in the writing process

During the preparation of this work the author(s) used ChatGPT to improve readability of a few sections of the manuscript. After using this tool/service, the author(s) reviewed and edited the content as needed and take(s) full responsibility for the content of the publication.

## Notes

### Competing Interest Statement

The authors have declared no competing interest.

