## Supplementary material for "Coupling renewable energy infrastructure planning and biodiversity conservation: a modeling-based framework": Suplementary material

**--**

**Supplementary materials**

Jérémy S.P. Froidevaux^1,2*^, Isabelle Le Viol^2^, Kévin Barré^2^, Yves Bas^2,3,^, Christian Kerbiriou^2^

^1^ University of Stirling, Biological and Environmental Sciences, Faculty of Natural Sciences, FK9 4LA Stirling, UK.

^2^ Centre d'Ecologie et des Sciences de la Conservation (CESCO), Muséum national d'Histoire naturelle, Centre National de la Recherche Scientifique, Sorbonne Université, Station Marine, 29900 Concarneau / 75005 Paris, France.

^4^ CEFE, Université de Montpellier, CNRS, EPHE, IRD, 34090 Montpellier, France.

**Supplement S1. Location of the field sites acoustically sampled through the Vigie-Chiro program for the period 2015-2020**

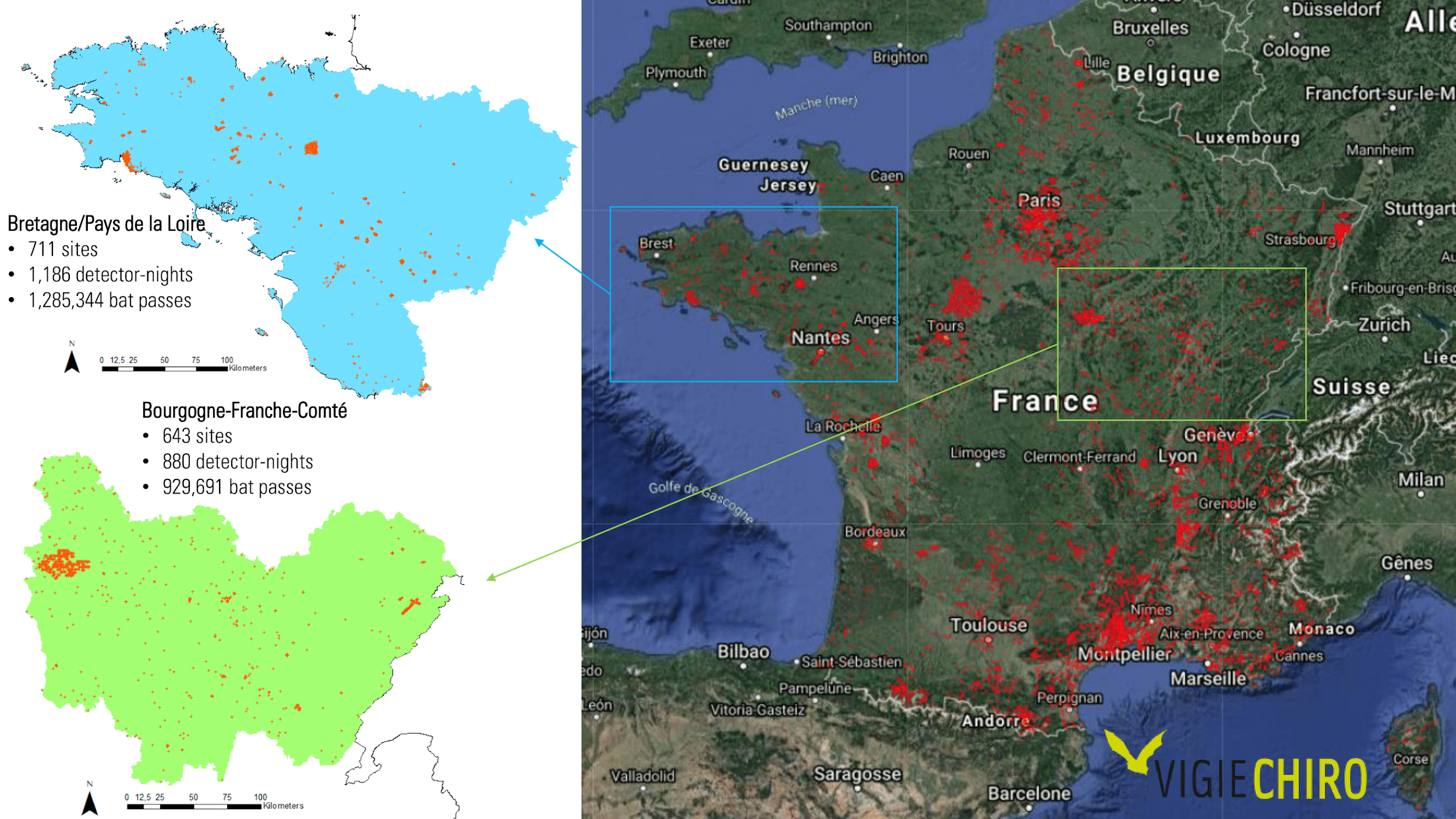

**Figure S1-1.** Maps displaying the field sites (red dots) that were acoustically sampled through the French national-scale citizen-science bat monitoring program “Vigie-Chiro” for the period 2015-2020 in France (right panel) and more specifically in the region Bourgogne-Franche-Comté (BFC; green area) and the regions Bretagne and Pays de la Loire (BPL; blue area) (left panel).

**Supplement S2. Land cover composition around Vigie-Chiro sites**

**Representativeness of land cover around Vigie-Chiro sites**

To check whether land cover around the Vigie-Chiro sites surveyed were representative of the overall land cover of the two study areas (Bourgogne-Franche-Comté BFC and Bretagne-Pays de la Loire BPL) - in other words to check for spatial bias in citizen-science data (Newson et al. 2015, 2017) - we first compared the amount of dominant land cover around (i) Vigie-Chiro sites (N_BFC_ = 643; N_BPL_ = 711) and (ii) centroids of a 0.5 x 0.5 km systematic grid covering the study areas (N_BFC_ = 193,848; N_BPL_ = 244,099). To do so, we created a buffer of 0.25 km radius around each Vigie-Chiro site and each grid centroid, and extracted the amount of the dominant land cover, namely: cropland, deciduous forest, grassland and urban area. Barplots in Figure S2-1 indicated that Vigie-Chiro sites were representative of the main land cover of the study areas. Only deciduous forests were slightly over-sampled in BFC. We then investigated specific land cover combinations i.e. gradient of a given land use cover within three classes of a different land use cover and distance to the nearest water source. We also found that Vigie-Chiro sites were representative of the combinations occurring in the study areas (e.g. Figures S2-2 and S2-3).

**Potential biases in land cover representativeness between bat detectors**

Even though detectors were configured to limit heterogeneity in bat detection probabilities among detector types, residual heterogeneity may still occur and bias the results if detector type is confounded with habitat type surveyed (e.g. if only one detector type is used to survey a given habitat). We therefore visually checked the extent to which the types of detectors used was related to habitat composition of Vigie-Chiro sites at 250 m radius buffer scale; we found no apparent confounding effect (e.g. Figure S2-4).

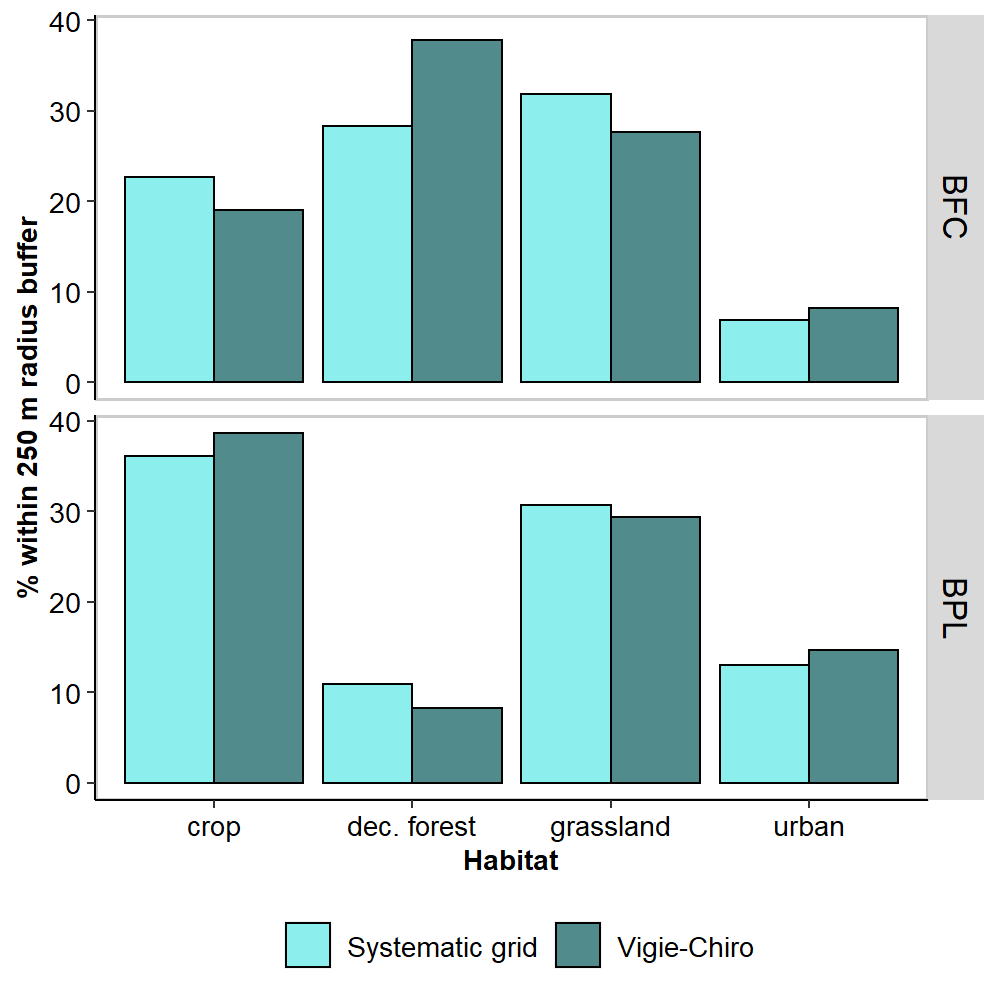

**Figure S2-1**. Percentage of dominant land cover type within 250m radius buffer around Vigie-Chiro sites and centroids of a 0.5 x 0.5 km grid in the two study areas (Bourgogne-Franche-Comté BFC and Bretagne-Pays de la Loire BPL).

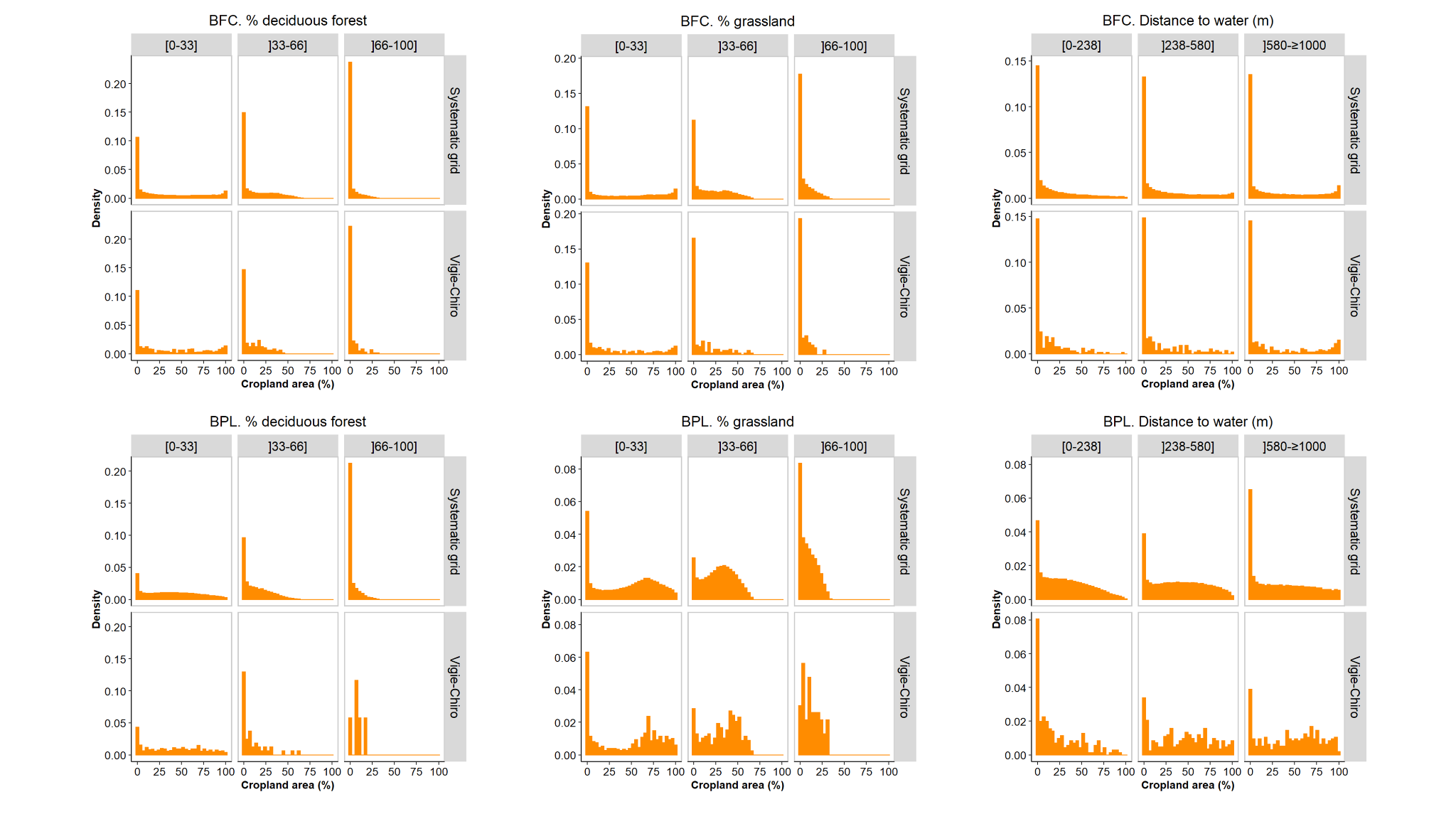

**Figure S2-2**. Percentage of cropland area within 250m radius buffer around Vigie-Chiro sites and centroids of a 0.5 x 0.5 km grid in the two study areas (Bourgogne-Franche-Comté BFC and Bretagne-Pays de la Loire BPL) within different classes (bounded by percentile 33%, 66% and 100%) of (i) amount of deciduous forest, (ii) amount of grassland, and (iii) distance to water.

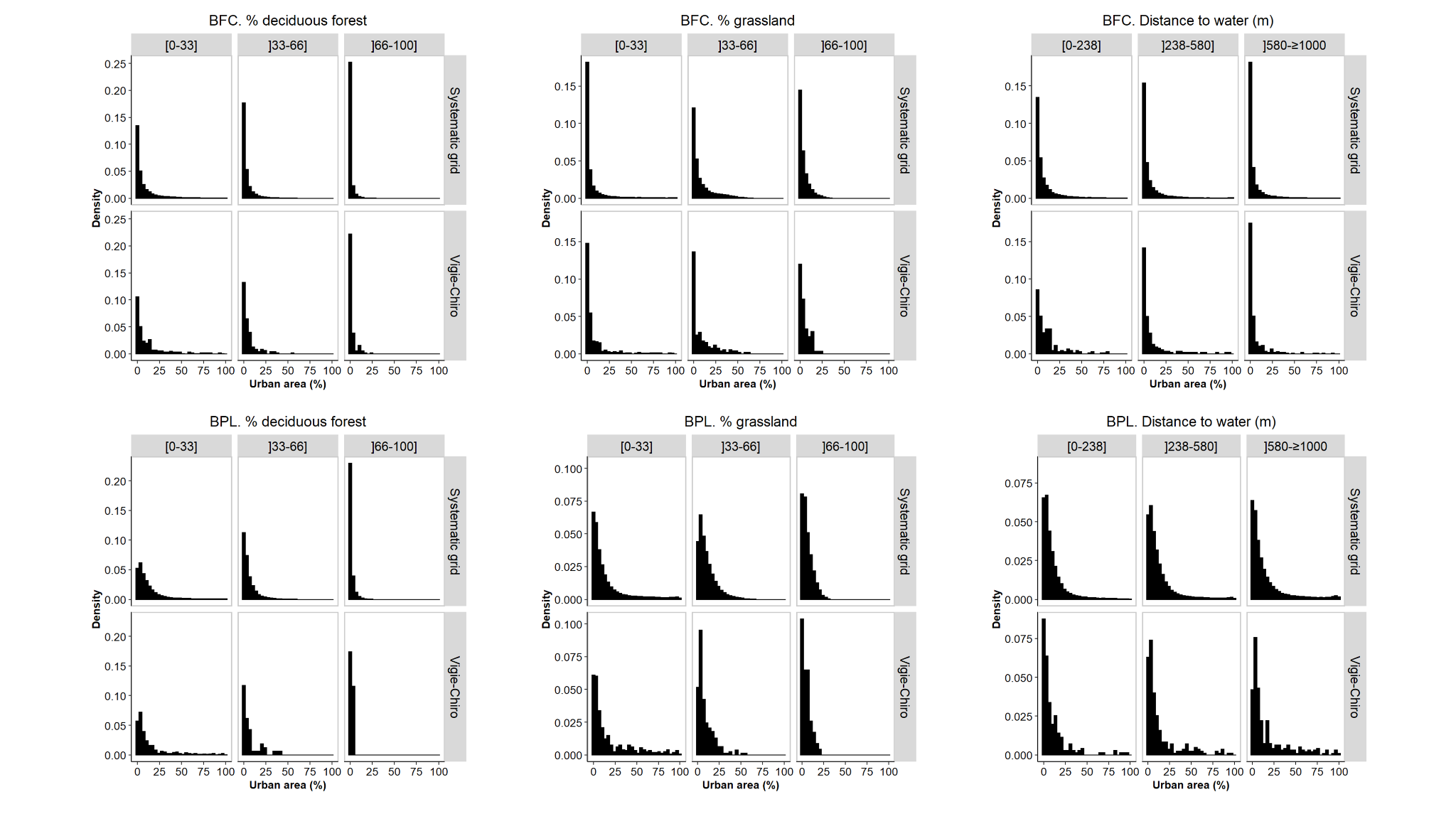

**Figure S2-3**. Percentage of urban area within 250m radius buffer around Vigie-Chiro sites and centroids of a 0.5 x 0.5 km grid in the two study areas (Bourgogne-Franche-Comté BFC and Bretagne-Pays de la Loire BPL) within different classes (bounded by percentile 33%, 66% and 100%) of (i) amount of deciduous forest, (ii) amount of grassland, and (iii) distance to water.

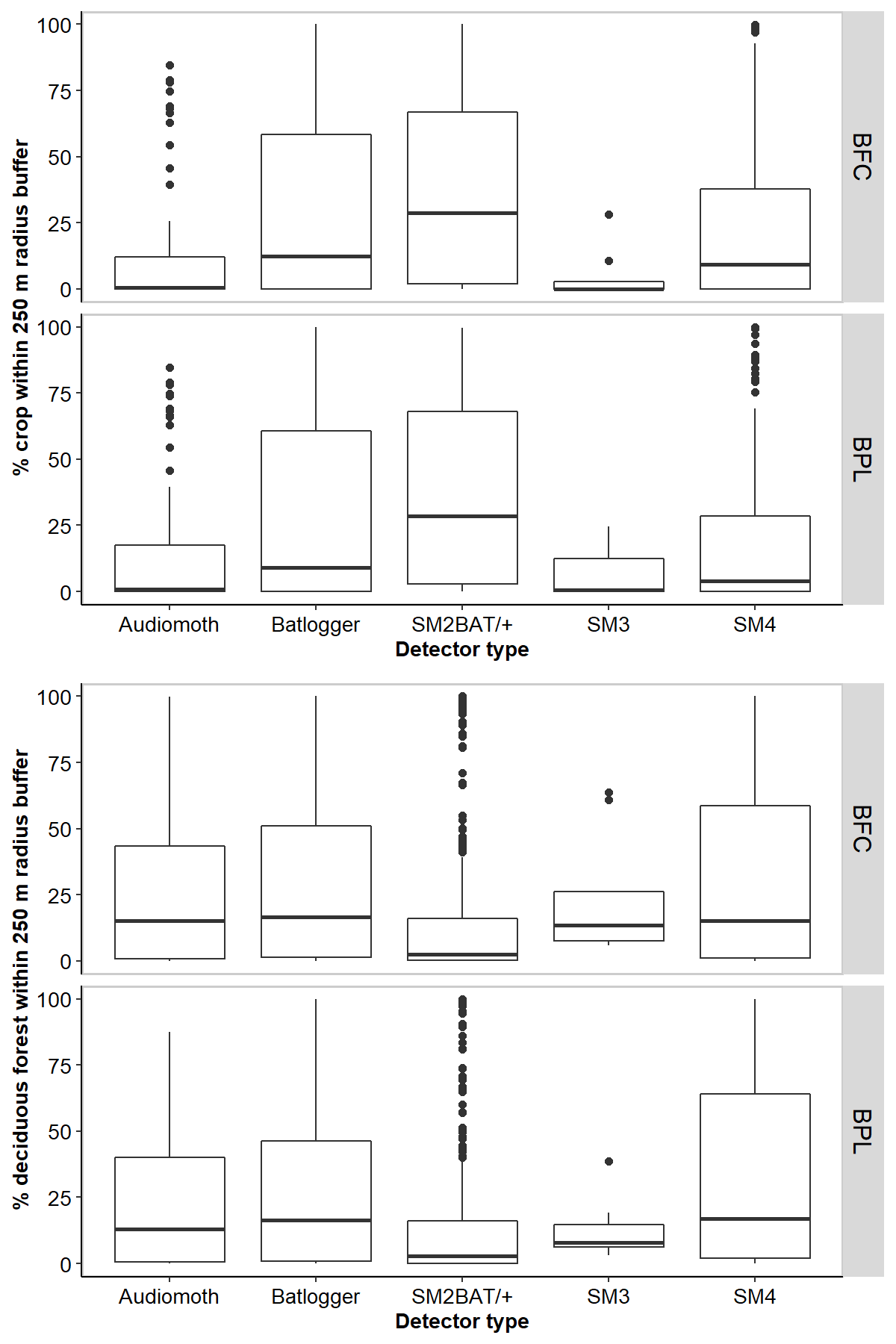

**Figure S2-4**. Percentage of the two dominant land use types (cropland and deciduous forest) within 250m radius buffer around Vigie-Chiro sites depending on the detector type used in the two study areas (Bourgogne-Franche-Comté BFC and Bretagne-Pays de la Loire BPL).

**Supplement S3. Sources, temporal coverage, and resolution of the model predictors**

**Table 3-1.** List of the predictors with their source and their temporal coverage and resolution (when applicable) included in the random forest models.

| **Predictor** | **Source** | **Temporal coverage** | **Resolution** |
| --- | --- | --- | --- |
| Altitude (m a.s.l.) | IGN BD Alti  https://geoservices.ign.fr/documentation/diffusion/telechargement-donnees-libres.html#bd-alti | 2020 | 75 m |
| Slope (°) | IGN BD Alti  https://geoservices.ign.fr/documentation/diffusion/telechargement-donnees-libres.html#bd-alti | 2020 | 75 m |
| % of deciduous forests | CES OSO land cover data  https://osr-cesbio.ups-tlse.fr/~oso/ | 2018 | 10 m |
| Dist. to deciduous forests (m) | CES OSO land cover data  https://osr-cesbio.ups-tlse.fr/~oso/, | 2018 | 10 m |
| % Linear small woody features | Copernicus  https://land.copernicus.eu/pan-european/high-resolution-layers/small-woody-features | 2014-2016 | 5 m |
| Dist. to linear small woody features (m) | Copernicus  https://land.copernicus.eu/pan-european/high-resolution-layers/small-woody-features | 2014-2016 | 5 m |
| Density of rivers (m/ha) | Eaufrance BD Carthage  https://www.data.gouv.fr/fr/datasets/cours-deau-metropole-2017-bd-carthage | 2017 | vector |
| Dist. to freshwater body (m) | Eaufrance BD Carthage  https://www.data.gouv.fr/fr/datasets/plans-deau-metropole-2017-bd-carthage | 2017 | vector |
| Edge density of major land-cover classes* | CES OSO land cover data  https://osr-cesbio.ups-tlse.fr/~oso/ | 2018 | 10 m |
| Shannon’s diversity index of major land-cover classes* | CES OSO land cover data  https://osr-cesbio.ups-tlse.fr/~oso/ | 2018 | 10 m |
| Artificial night-time light brightness | NOAA https://ngdc.noaa.gov/eog/viirs/download_dnb_composites.html | 2016 | 350 m |
| % of urban areas | CES OSO land cover data  https://osr-cesbio.ups-tlse.fr/~oso/ | 2018 | 10 m |
| Dist. to urban areas (m) | CES OSO land cover data  https://osr-cesbio.ups-tlse.fr/~oso/ | 2018 | 10 m |
| % of croplands | CES OSO land cover data  https://osr-cesbio.ups-tlse.fr/~oso/ | 2018 | 10 m |
| Dist. to croplands (m) | CES OSO land cover data  https://osr-cesbio.ups-tlse.fr/~oso/ | 2018 | 10 m |
| Density of major roads (m/ha) | IGN Route 500 https://geoservices.ign.fr/documentation/diffusion/telechargement-donnees-libres.html#route-500 | 2020 | vector |
| Dist. to major roads (m) | IGN Route 500 https://geoservices.ign.fr/documentation/diffusion/telechargement-donnees-libres.html#route-500 | 2000-2020 | vector |
| Density of operational wind turbines (no/ha) | Bretagne: https://geobretagne.fr/mapfishapp/  Pays de la Loire: https://carto.sigloire.fr/  Bourgogne-Franche-Comté: https://cartes.ternum-bfc.fr/ | 2000-2020 | vector |
| Dist. to operational wind turbines (m) | Bretagne: https://geobretagne.fr/mapfishapp/  Pays de la Loire: https://carto.sigloire.fr/  Bourgogne-Franche-Comté: https://cartes.ternum-bfc.fr/ | 2000-2020 | vector |
| Human population density (no/km^2^) | GHSL  https://ghsl.jrc.ec.europa.eu/ghs_pop.php | 2015 | 250 m |
| Quietness suitability index | EEA  https://www.eea.europa.eu/data-and-maps/figures/quietness-suitability-index-qsi-2 | 2016 | 100 m |
| Bioclimatic variables | CHELSEA database v1.2  Karger et al. 2017a; Karger et al. 2017b | 1979-2013. | 30 arc sec (~1 km) |

*Calculated in R with *landscapemetrics* package (Hesselbarth et al. 2019)

**Supplement S4. Percentage of variance explained by the models**

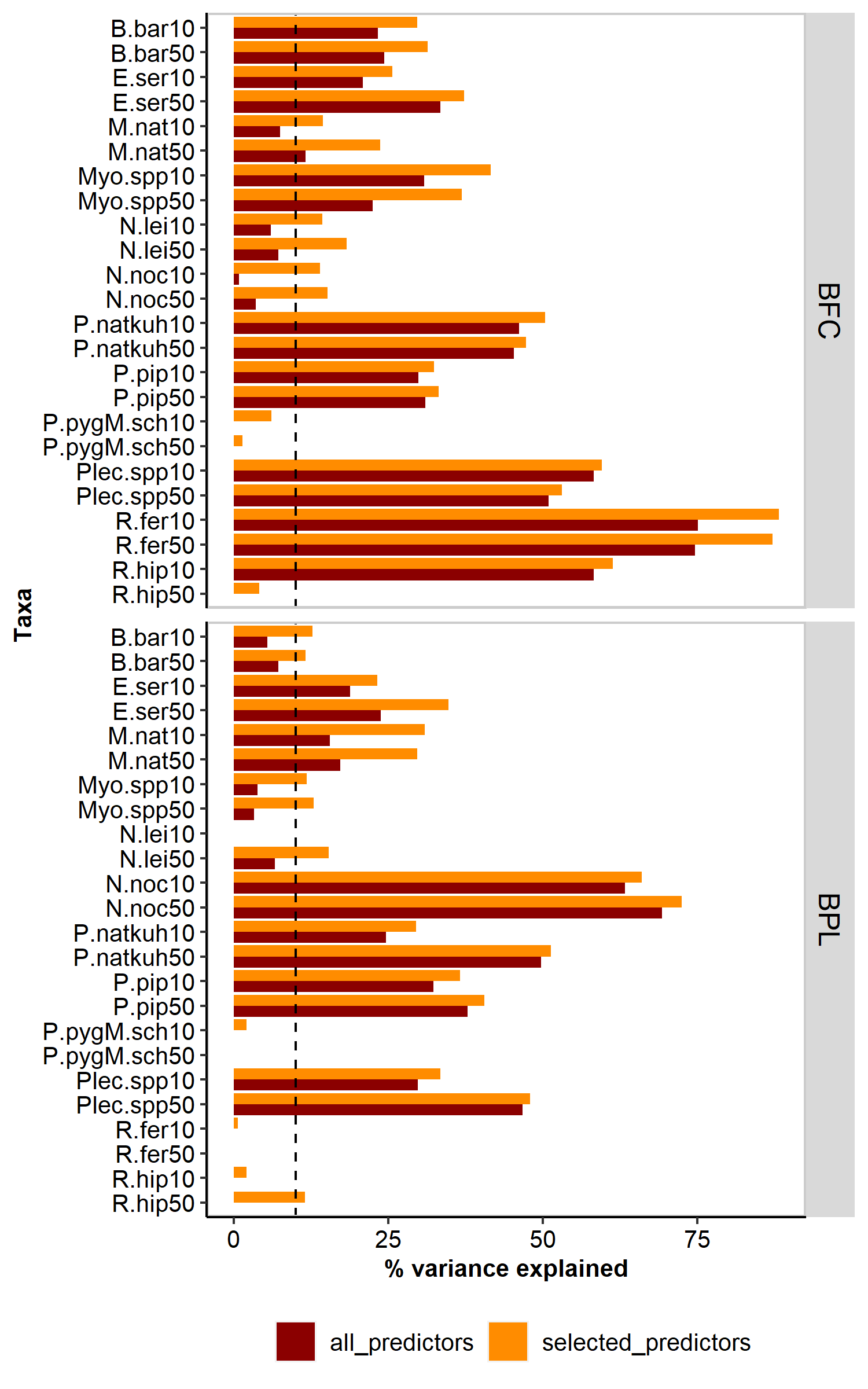

**Figure S4-1**. Bar plots depicting the percentage of variance explained (here referred to as a measure of how well the out-of-bag (OOB) predictions capture the variability of the target variables in the training set) extracted for each random forest model - i.e. one per taxa and per area (BFC: Bourgogne-Franche-Comté, BPL: Bretagne-Pays de la Loire) and considering two acoustic datasets (10 vs 50% maximum error risk tolerance). Red bars depict the percentage of variance explained for models that included all predictors while orange bars depict the percentage of variance explained for models that included only the predictors retained after model selection. Note that only one spatial dataset (the full one) was used in this step.

**Supplement S5. ‘Overview, Data, Model, Assessment and Prediction’ protocol**

### Overview

##### Authorship

Contact :

##### Model objective

Model objective: Mapping and interpolation

Target output: Activity

##### Focal Taxon

Focal Taxon: Bat

##### Location

Location: Bourgogne-Franche Comté and Bretagne-Pays-de-la-Loire, France

##### Scale of Analysis

Spatial extent: -4.79640, 7.14331, 46.2965445, 48.87011 (xmin, xmax, ymin, ymax)

Spatial resolution: Min: 0.005; Max: 0,350

Temporal extent: 2015-2020

Temporal resolution: Day

Boundary: political

##### Biodiversity data

Observation type: citizen science, standardised monitoring data, field survey

Response data type: counts

##### Predictors

Predictor types: climatic, habitat, topographic, anthropogenic

##### Hypotheses

Hypotheses: We hypothesized that bats respond to environmental variables and anthropogenic stressors at different spatial scales, from local to landscape scales.

##### Algorithms

Modelling techniques: randomForest

<Model complexity>

Model averaging: NA

##### Workflow

Model workflow: See Figure 1 in the main text for a more detailed description of the modeling-based framework.

Modelling steps:

##### Software

Software: Software: R v4.2.1. Packages: -randomForest 4.7-1.1 -VSURF 1.1.0

Code availability: Zenodo (< add link here >)

Data availability: Zenodo (< add link here >)

### Data

##### Biodiversity data

Taxon names: *Barbastella barbastellus*, *Eptesicus serotinus*, *Myotis nattereri*, *Myotis* spp. (excluding *M. nattereri*), *Nyctalus leisleri*, *Nyctalus noctula*, *Pipistrellus* *pipistrellus*, *Pipistrellus* *nathusii*/*kuhlii*, *Pipistrellus* *pygmaeus*/*Miniopterus* *schreibersii*, *Plecotus* spp., *Rhinolophus* *ferrumequinum*, *Rhinolophus* *hipposideros*.

Taxonomic reference system: Handbook of the mammals of Europe. Russo, D. Chiroptera. (Springer Cham, 2023)

Ecological level: species

Data sources: Data source: <https://www.vigienature.fr/fr/chauves-souris> and <https://vigiechiro.herokuapp.com/#/accueil> Accession date: Nov 2020

Sampling design: The French national-scale citizen-science bat monitoring program ‘Vigie-Chiro’ has been created and coordinated by the French National Museum of Natural History since 2006. There are three main acoustic monitoring-based protocols proposed to volunteers: the car transect protocol, the walk transect protocol and the stationary point protocol. Hereafter, only data from the stationary point protocol were considered. For this protocol, volunteers set stationary recording devices to record sounds between 8 and 192 kHz throughout the entire night, from 30 min before sunset to 30 min after sunrise (Dubos et al. 2021). The Vigie-Chiro program was originally designed to study bat population trends in France. One major prerequisite to meet such an objective is to ensure that the proportion of habitats sampled is largely representative of French land-use. Thus, volunteers taking part in the stationary point protocol are encouraged to survey randomly selected sampling points, even though this protocol also offers flexibility for volunteers to focus on particular sites. Volunteers are invited to geolocate their device in a systematic national 2-km square grid with several options: (i) volunteers can choose the square monitored or (ii) used a randomly chosen square in a radius of 10km from their home, (i.e. on average one square randomly chosen between 80 possible squares). In addition, within each square volunteers can (i) choose the site monitored or (ii) used a predefined site. Regardless of the option chosen, sampling points should be located at least 200 meters from each other. Preliminary analysis showed that Vigie-Chiro sites covered similar land-use type gradients as sites randomly selected in France (see Mariton et al. 2022). Volunteers may use a variety of full-spectrum ultrasound recorders, including Song Meter SM2BAT+, SM3BAT, SM4BAT, SMminiBAT (Wildlife Acoustics Inc., Concord, MA, USA), Batlogger (Elekon, Luzern, Switzerland), Passive Recorder (<https://framagit.org/PiBatRecorderPojects/TeensyRecorders>), Anabat Swift (Titley Scientific, Brendale, Australia) and AudioMoth (Hill et al. 2018), as long as the device settings meet the Vigie-Chiro criteria (e.g. standardised trigger setting to limit heterogeneity in detectability). Sampling sessions occurred whenever possible when weather conditions were favourable, i.e. no rain, temperature above seasonal normal and avoiding strong winds (<30 km.h-1), even though long-term monitoring over consecutive nights did not always meet these criteria. Volunteers are encouraged to monitor bats at least twice a year: a first visit during June and July, when females are expected to give birth and feed their offspring; and a second visit between 15 August and 31 September, when juveniles are flying and migratory adults are expected to be contacted.

Sample size: See table 1 in the main text.

Clipping: Mask: Bourgogne-Franche Comté and Bretagne-Pays-de-la-Loire regions, France

Scaling: NA

Cleaning: Filtering steps: We restricted our selection to sites (i) sampled between May and October (i.e. period of highest bat activity), (ii) with microphones placed at <5 m height, (iii) located away (>200 m) from known roosts, and (iv) at lower altitude (<800 m a.s.l.) to avoid excessive heterogeneity due to mountainous environmental characteristics.

Absence data: NA

Background data: NA

Errors and biases: (1) Potential error in bat echolocation call identification: We followed the method proposed by Barré et al. (2019) to account for potential automated identification errors, i.e. we used the confidence index to retain two separate datasets: (i) one dataset of bat passes with maximum error risk tolerance of 10%; and (ii) another dataset of bat passes with maximum error risk tolerance of 50%. The first threshold is cautious, aiming to minimize false positives, while the second one is less conservative, allowing for a larger amount of data to be retained. Bat activity per night corresponded to the sum of bat passes recorded. (2) Potential spatial bias: Because some Vigie-Chiro sites were spatially clustered (especially in Bretagne-Pays-de-la-Loire) - thus potentially leading to an overestimation of model performance - we ran the final models using two datasets that included either all the sites or only spatially independent ones, i.e. a subset of the full dataset containing only sites that were located >500 m away from each other. Please see supplementary material S2 for a thorough assessment of potential bias with regards to land cover composition around Vigie-Chiro sites.

References: Barré, K., Le Viol, I., Julliard, R., Pauwels, J., Newson, S.E., Julien, J.F., Claireau, F., Kerbiriou, C., Bas, Y., 2019. Accounting for automated identification errors in acoustic surveys. Methods in Ecology and Evolution 10, 1171-1188.

##### Data partitioning

Training data: The activity dataset for each target species was partitioned by randomly splitting into 80% for model training and 20% for model evaluation.

Validation data: We iterated 500 times the fivefold cross‐validations.

##### Predictor variables

Predictor variables: Full details are provided in Supplement S3.

Data sources: Full details are provided in Supplement S3.

### Model

##### Multicollinearity

Multicollinearity: See section 'Model selection strategy'.

##### Model settings

randomForest: ntree (500), mtry (1), maxnodes (maximum possible (subject to limits by nodesize))

Model settings (extrapolation): Not relevant

##### Model estimates

We checked variable importance with the mean decrease accuracy.

##### Model selection - model averaging - ensembles

Model selection: While Random Forest can operate with large numbers of variables and is largely insensitive to multicollinearity, it is recommended to proceed to variable selection to improve overall model performance. We therefore conducted a variable selection procedure using the R-package vsurf and retained the smaller set of variables sufficient for prediction purposes. In this process, predictor variables were assessed for their individual impact on model performance and the resulting list of selected predictors was refined by eliminating redundancy.

##### Analysis and Correction of non-independence

Spatial autocorrelation: Not statistically assessed. See section 'Details on potential errors and biases in data' for the alternative approach used.

Temporal autocorrelation: Not statistically assessed. Julian day was added as a predictor.

Nested data: We used participant ID and site ID as strata to account for the stratified structure of the citizen-science data.

##### Threshold selection

Threshold selection: Not relevant

### Assessment

##### Performance statistics

Performance on training data: Percentage of variance explained

Performance on validation data: Normalized Root Mean Square Error statistic

##### Plausibility check

Expert judgement: Manual check of predicted bat activity in a subset of random points and wind turbines approved for construction.

### Prediction

##### Prediction output

Prediction unit: Bat activity (number of bat passes per night)

##### Uncertainty quantification

Scenario uncertainty: Not relevant.

Novel environments: Not assessed.

**Supplement S6. Model predictive performance**

Model predictive performance varied with respect to species, area and spatial dataset considered (Figure S6-1). We observed very little differences in predictive performance between the two acoustic datasets used. Species-specific variations in model predictive performance were expected as it has been suggested that species with low abundance and frequency of occurrence are more difficult to predict (Waldock et al. 2022) and that predictive performance may depend on species traits (Luan et al. 2020). Thus, variations in bat occurrence and activity across BFC and BPL, which have contrasting past and current land-use and climate, may also explain the differences observed in performance statistic values between the two study areas. When restricting the dataset to spatially independent sites (here sites >500 m from each other), model predictive performance of all models was reduced regardless of the study areas and the acoustic datasets. Nevertheless, reduction in predictive performance was more marked in BPL where sampling sites tended to be clustered.

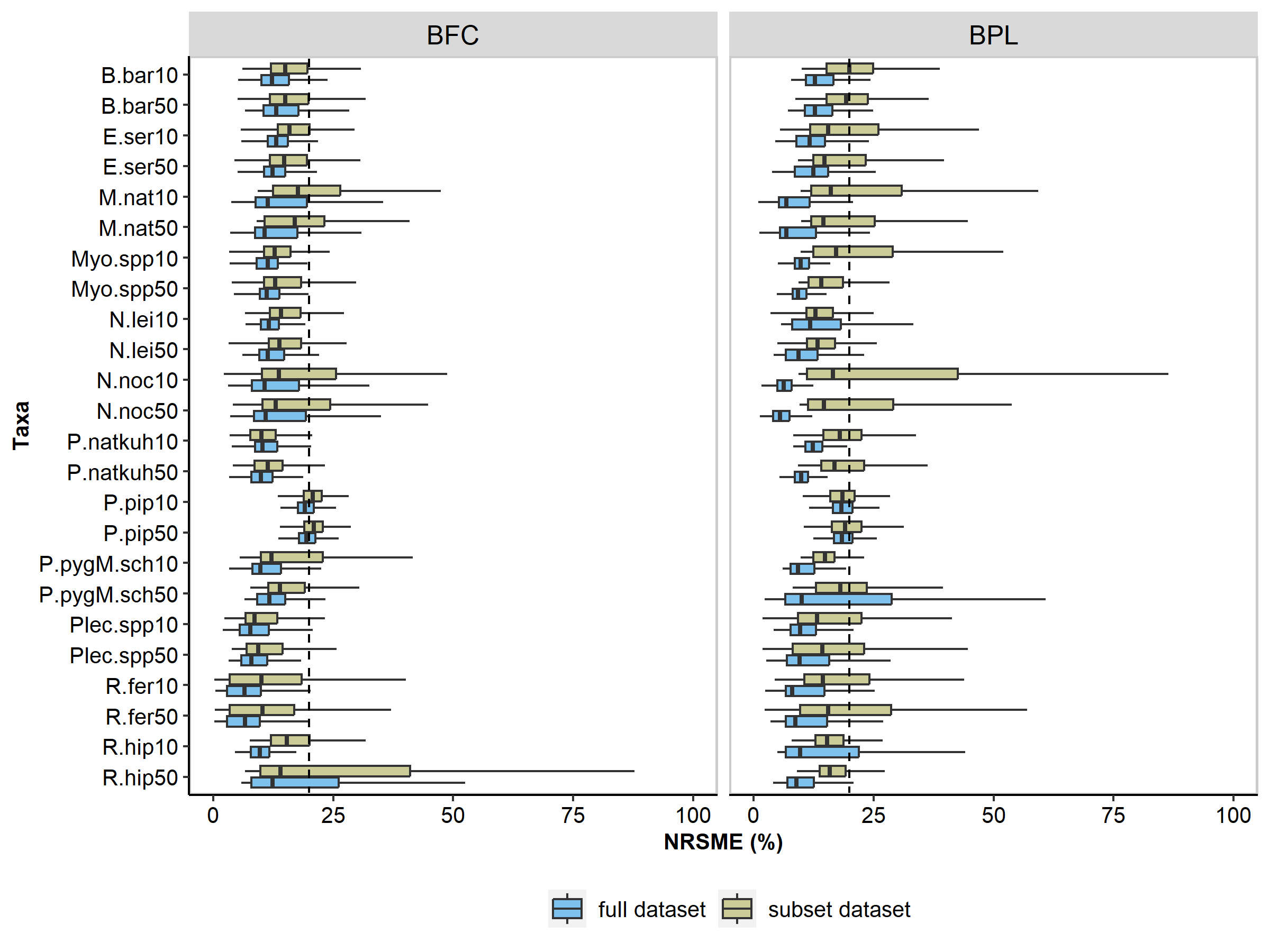

**Figure S6-1**. Boxplot of Normalized Root Mean Square Error statistic derived from a fivefold cross‐validation procedure iterated 500 times to assess predictive performance of the random forest models. The boxplots display the interquartile range box (top line = 75% of the data ≤ this value; middle line = median; lower line = 25% of the data ≤ this value) and the lower and upper whiskers (minimum and maximum data points). Lower NRMSE values indicate higher performance, and the dashed line represents the 20% threshold. NRMSE were calculated for each final random forest, i.e. one per taxa and per area (BFC: Bourgogne-Franche-Comté, BPL: Bretagne-Pays de la Loire) and considering two acoustic datasets (10 vs 50% maximum error risk tolerance) and two spatial datasets (full vs subset dataset with spatially independent sites).
